# p300 KAT regulates SOX10 stability and function in human melanoma

**DOI:** 10.1101/2024.02.20.581224

**Authors:** Aaron Waddell, Nicole Grbic, Kassidy Leibowitz, W. Austin Wyant, Sabah Choudhury, Kihyun Park, Marianne Collard, Philip A. Cole, Rhoda M. Alani

## Abstract

SOX10 is a lineage-specific transcription factor critical for melanoma tumor growth, while SOX10 loss-of-function drives the emergence of therapy-resistant, invasive melanoma phenotypes. A major challenge has been developing therapeutic strategies targeting SOX10’s role in melanoma proliferation, while preventing a concomitant increase in tumor cell invasion. Here, we report that the lysine acetyltransferase (KAT) *EP300* and *SOX10* gene loci on Chromosome 22 are frequently co-amplified in melanomas, including UV-associated and acral tumors. We further show that p300 KAT activity mediates SOX10 protein stability and that the p300 inhibitor, A-485, downregulates SOX10 protein levels in melanoma cells via proteasome-mediated degradation. Additionally, A-485 potently inhibits proliferation of SOX10+ melanoma cells while decreasing invasion in AXL^high^/MITF^low^ melanoma cells through downregulation of metastasis-related genes. We conclude that the SOX10/p300 axis is critical to melanoma growth and invasion, and that inhibition of p300 KAT activity through A-485 may be a worthwhile therapeutic approach for SOX10-reliant tumors.

## Introduction

Treatment for advanced melanoma has been revolutionized in the last decade through the development of targeted (MAPK inhibitor) therapies and immunotherapies. ^1^ While these therapeutic strategies offer significant short-term clinical benefits, low overall response rates, acquired therapy resistance and a lack of additional targetable mutations suggest the need for new therapeutic strategies to significantly impact melanoma cure rates. ^1^

Melanoma biology is characterized by phenotype plasticity, which is driven by selective pressures and associated with reversible transcriptional programs controlled by master regulator transcription factors including MITF, ZEB1/ZEB2 and SOX10. ^2,3^ SOX10 is a neural crest lineage-specific transcription factor that is highly expressed in the melanocytic, transitory and neural crest-like melanoma phenotypes^4^ and is essential for melanoma development^5–7^, rapid growth^5,7,8^, and regulation of tumor immunogenicity. ^9–12^ Loss of SOX10 induces an undifferentiated melanoma phenotype characterized by slow-growth, increased metastatic potential and MAPK-inhibitor (MAPKi) resistance. ^8,13^ While key regulators of SOX10 function are under intense investigation and may serve as potential therapeutic strategies to target SOX10 tumorigenic activity in melanoma^14–25^, such studies have failed to yield meaningful progress to date.

We have previously reported that genetic knockdown (KD) of the transcriptional coactivator p300 in melanoma cells results in significantly lower expression of SOX10 and MITF proteins. ^26^ P300 and its close paralog CBP are well-characterized lysine acetyltransferases (KATs) ^27,28^ and p300 has been shown to promote melanoma progression through multiple mechanisms including regulation of MITF function^26,29^, direct acetylation of BRAF to enhance its kinase activity^30^, an emerging role in therapy resistance^31^ and as a potential driver of acral melanoma through genetic amplification of the *EP300* locus. ^32^ Importantly, we and others have previously demonstrated that melanoma cells with high MITF expression are particularly sensitive to growth inhibition by the p300/CBP KAT inhibitor A-485. ^26,33,34^ Given the impact of p300 activity on melanoma growth, direct regulation of MITF transcription by p300, and the concomitant loss of SOX10 and MITF protein expression following p300 inhibition in melanoma cells, we sought to further clarify the connection between SOX10, MITF and p300.

Here we report that *SOX10* and *EP300* are commonly co-amplified in both UV-associated cutaneous and acral melanomas. Strikingly, we find that p300 KAT activity is essential for SOX10 protein stability and that inhibition of p300 KAT with the small molecule inhibitor, A-485, inhibits the expression of SOX10 target genes and SOX10-dependent melanoma growth in SOX10+ melanoma cell lines, regardless of MITF status. Notably, we find that p300 KAT activity is essential for the emergence of the invasive phenotype upon SOX10-loss in human melanomas, and that A-485 is capable of inhibiting both tumor cell proliferation and invasion in melanoma cells as a single agent. These studies identify p300 as a novel regulator of tumor phenotypes in melanoma through promotion of SOX10 function and as a key epigenetic mediator of the invasive melanoma phenotype in the absence of SOX10. We therefore suggest that *EP300/SOX10* co-amplification in melanoma may serve as a biomarker for tumors which would benefit from therapeutic p300 inhibition.

## Materials and Methods

### EP300 and SOX10 copy number and expression data for patients and cell lines

For patient data, EP300 and SOX10 copy number segment data was obtained for the TCGA SKCM, MSKCC (2014), MSKCC (2017), DFCI (2015) melanoma datasets utilizing the cBioPortal service (https://www.cbioportal.org/). Copy number segment data was analyzed via GISTIC2.0 on default settings utilizing the Gene Pattern (https://www.genepattern.org) online service to determine the copy number status of patient samples. Acral melanoma samples from multiple datasets were combined (Combined Acral) to achieve a higher sample size for determination of copy number status. mRNA expression data for EP300 and SOX10 were obtained through cBioPortal for the TCGA SKCM dataset and correlated with copy number status. For cell line data, EP300 and SOX10 relative copy number, absolute copy number and mRNA expression data was obtained from the Cancer Dependency Map (depmap.org) and plotted as indicated in the figures.

### Cell culture

Melanoma cell lines 451Lu, Sk-Mel-28, Sk-Mel-24, WM983B, 1205Lu, WM793 and A375 were obtained from Dr. Meenard Herlyn (The Wistar Institute, Philadelphia, PA). SK-MEL-30 and IPC-298 cells were obtained from Dr. Anurag Singh (Boston University, Boston, MA, USA). Acral melanoma cell line CO79 was obtained from Dr. Nick Hayward (Berghofer Medical Research Institute, Herston, Australia). YUSEEP and YUHIMO were obtained from Dr. Ruth Halaban (Yale University, New Haven, Connecticut, USA). Sk-Mel-28 BRAFi-R cells were obtained from Dr. Deborah Lang (Boston University, Boston, MA). 451Lu BRAFi-R and A375 BRAFi-R cells were obtained from Dr. Jong-In Park (Medical College of Wisconsin, Wauwatosa, WI). All cell lines were authenticated in the last two years by LabCorp (Burlington, NC, USA) and were confirmed mycoplasma free before experimentation via DAPI staining. Cell lines were grown at 37 °C with 5% CO2 in a humidified atmosphere. All cell lines were cultured in Dulbecco’s Modified Eagle Medium (Gibco, ThermoFisher Scientific, Grand Island, NY, USA) supplemented with 10% bovine calf serum (Gibco, ThermoFisher Scientific, Grand Island, NY, USA), 1% penicillin/streptomycin (Gibco, ThermoFisher Scientific, Grand Island, NY, USA), and 2mM L-glutamine (Gibco, ThermoFisher Scientific, Grand Island, NY, USA) (designated as complete DMEM media). BRAFi-R cells were continuously cultured in complete DMEM media containing 5 µM PLX4032 (MedChemExpress Monmouth Junction, NJ, USA).

### Compounds

A-485, Cycloheximide (CHX), MG-132 and CCS1477 were purchased from MedChemExpress Monmouth Junction, NJ, USA).

### Immunoblotting

For immunoblots, the indicated cell lines seeded in 6-well plates in complete media. When cells reached 90% confluency, they were treated with DMSO or A-485 (5 µM) for 24 hours for analysis of SOX10, MITF and DCT protein levels. For analysis of SOX10 protein stability, cells were treated with Cycloheximide (50 µg/mL) or Cyclohexamide (50 µg/mL) + MG-132 (5 µM) for the indicated time points. For analysis of SOX10 degradation by A-485, cells were treated with DMSO, A-485 (5 µM), MG-132 (5 µM) or A-485 + MG-132 (5 µM) for 16 hours. After the indicated drug treatment period, cells were washed with PBS, the PBS was aspirated and cells were lysed in complete M-PER buffer (Thermo Scientific, Rockford, IL, USA). Complete M-PER buffer was prepared through addition of Protease & Phosphatase Inhibitor Cocktail (1X final) (Thermo Scientific, Rockford, IL, USA) to the M-PER buffer. The protein content of the resulting lysate was quantified using the BCA Protein Assay Kit (Thermo Scientific, Rockford, IL, USA) according to manufacturer instructions. 30 µg of protein lysate were then run per lane on a 10% polyacrylamide gel (Bio-Rad Laboratories, Hercules, CA, USA) and transferred to a PVDF membrane (Bio-Rad Laboratories, Hercules, CA, USA) using the Trans-Blot® Turbo™ Transfer System (Bio-Rad Laboratories, Hercules, CA, USA) according to manufacturer instructions. Membranes were blocked with 1X TBST (Sigma-Aldrich, St Louis, MO, USA) containing 5% non-fat milk (Sigma-Aldrich, St Louis, MO, USA). Membranes were then incubated overnight with primary antibodies in 1X TBST containing 5% BSA (Sigma-Aldrich, St Louis, MO, USA). Membranes with multiple probes were stripped using 0.2 M NaOH before probing with the next primary antibody. Primary antibodies: SOX10 (SC-365692, Santa Cruz Biotechnology, Dallas, TX, USA), SOX10 (69661, Cell Signaling Technology, Danvers, MA, USA), MITF (12590, Cell Signaling Technology, Danvers, MA, USA), DCT (SC-74439, Santa Cruz Biotechnology, Dallas, TX, USA), GAPDH (SC-365062, Santa Cruz Biotechnology, Dallas, TX, USA). Secondary antibodies: anti-mouse IgG HRP (7076, Cell Signaling Technology, Danvers, MA, USA) and anti-rabbit (G21234, Invitrogen, ThermoFisher Scientific, Carlsbad, CA, USA). Immunoblots were visualized using the Chemiluminescent HRP Substrate kit (MilliporeSigma, Burlington, MA, USA) and the ChemiDoc™ XRS+ Molecular Imager® (Bio-Rad Laboratories, Hercules, CA, USA). Densitometry quantification of immunoblot bands was performed using ImageJ.

### RT-qPCR

The indicated cell lines were seeded in 6-well plates in complete media. When cells reached 90% confluency, they were treated with DMSO or A-485 (5 µM) for 24 hours. RNA was then extracted using the RNeasy (Qiagen, Germantown, MD, USA) kit according to manufacturer instructions. cDNA was then synthesized using the SuperScript™ III Reverse Transcriptase kit (Invitrogen, ThermoFisher Scientific, Carlsbad, CA, USA) according to manufacturer instructions. qPCR was performed on the Applied Biosystems Step One Plus Real Time PCR System according to the manufacturer instructions using the iQ SYBR® Green Supermix (Bio-Rad Laboratories, Hercules, CA, USA) kit. Primer sequences as follows: *18S* Forward: CTACCACATCCAAGGAAGCA, *18S* Reverse: TTTTTCGTCACTACCTCCCCG, *MITF* Forward: GGAAATCTTGGGCTTGATGGA, *MITF* Reverse: CCCGAGACAGGCAACGTATT, *DCT* Forward: TATTAGGACCAGGACGCCCC, *DCT* Reverse: TGGTACCGGTGCCAGGTAAC, *SOX10* Forward 1: CTTTCTTGTGCTGCATACGG, *SOX10* Reverse 1: AGCTCAGCAAGACGCTGG, *SOX10* Forward 2: TACCCGCACCTGCACAAC, *SOX10* Reverse 2: TTCAGCAGCCTCCAGAGC, *FN1* Forward: GCCGAATGTAGGACAAGAAGC, *FN1* Reverse: TGCCTCCACTATGACGTTGT, *THBS1* Forward: CCAGATGAACGGGAAACCCT, *THBS1* Reverse: CCTCCACAGGTGACAGAACAG, *MMP1* Forward: GGGCCACTATTTCTCCGCTT, *MMP1* Reverse: AAGGCCAGTATGCACAGCTT, *LAMC2* Forward: TACAGAGCTGGAAGGCAGGATG, *LAMC2* Reverse: GTTCTCTTGGCTCCTCACCTTG, *NT5E* Forward: AGTCCACTGGAGAGTTCCTGCA, *NT5E* Reverse: TGAGAGGGTCATAACTGGGCAC, *SFRP1* Forward: CAATGCCACCGAAGCCTCCAAG, *SFRP1* Reverse: CAAACTCGCTGGCACAGAGATG.

### RNA-sequencing

The indicated cell lines were seeded in 6-well plates in complete media. When cells reached 90% confluency, they were treated with DMSO or A-485 (5 µM) for 24 hours. RNA was then extracted using the RNeasy (Qiagen, Germantown, MD, USA) kit according to manufacturer instructions. RNA-sequencing was performed by Azenta Life Sciences. RNA library preparations, sequencing reactions, and bioinformatic analysis were conducted at GENEWIZ/Azenta Life Sciences LLC. (South Plainfield, NJ, USA).

### Library preparation and sequencing

RNA samples were quantified using Qubit 2.0 Fluorometer (Life Technologies, Carlsbad, CA, USA) and RNA integrity was checked using Agilent TapeStation 4200 (Agilent Technologies, Palo Alto, CA, USA). RNA sequencing libraries were prepared using the NEBNext Ultra II RNA Library Prep for Illumina using manufacturer’s instructions (NEB, Ipswich, MA, USA). Briefly, mRNAs were initially enriched with Oligod(T) beads. Enriched mRNAs were fragmented for 15 minutes at 94°C. First strand and second strand cDNA were subsequently synthesized. cDNA fragments were end repaired and adenylated at 3’ends, and universal adapters were ligated to cDNA fragments, followed by index addition and library enrichment by PCR with limited cycles. The sequencing libraries were validated on the Agilent TapeStation (Agilent Technologies, Palo Alto, CA, USA), and quantified by using Qubit 2.0 Fluorometer (Invitrogen, Carlsbad, CA) as well as by quantitative PCR (KAPA Biosystems, Wilmington, MA, USA). The sequencing libraries were clustered on a lane of a HiSeq flowcell. After clustering, the flowcell was loaded on the Illumina instrument (4000 or equivalent) according to manufacturer’s instructions. The samples were sequenced using a 2×150bp Paired End (PE) configuration. Image analysis and base calling were conducted by the Control software. Raw sequence data (.bcl files) generated the sequencer were converted into fastq files and de-multiplexed using Illumina’s bcl2fastq 2.17 software. One mismatch was allowed for index sequence identification. Raw sequence data (.bcl files) generated from Illumina HiSeq was converted into fastq files and de-multiplexed using Illumina’s bcl2fastq 2.20 software. One mis-match was allowed for index sequence identification.

### RNA-seq analysis

After investigating the quality of the raw data, sequence reads were trimmed to remove possible adapter sequences and nucleotides with poor quality using Trimmomatic v.0.36. The trimmed reads were mapped to the Homo sapiens GRCh38 reference genome available on ENSEMBL using the STAR aligner v.2.5.2b. Unique gene hit counts were calculated by using featureCounts from the Subread package v.1.5.2. Only unique reads that fell within exon regions were counted. After extraction of gene hit counts, the gene hit counts table was used for downstream differential expression analysis. Using DESeq2, a comparison of gene expression between groups of samples was performed. The Wald test was used to generate p-values and log2 fold changes. Genes with an adjusted p-value < 0.05 and absolute log2 fold change > 1 were called as differentially expressed genes for each comparison. Gene Ontology analysis was conducted using the top 500 (input limit = 500 genes) most differentially expressed genes using the GSEA Molecular Signatures Database (MSigDB) tool. ^46,60^

### Cell proliferation assays

For short-term (6-days) proliferation, the indicated cell lines were seeded in a 96-well plate in complete media. The cells were then cultured with the indicated compound for a total of 6-days before cell lysis and quantification of proliferation using the Quant-iT™ PicoGreen dsDNA Assay Kit (ThermoFisher Scientific, Grand Island, NY, USA). For long-term proliferation (12-days), the indicated cell lines were seeded in 6 cm plates in complete media. The cells were then cultured with the indicated compound for a total of 9 days and were passaged when they reached confluency. On the 9^th^ day of treatment, the cells were passaged and an equal number of cells per treatment were seeded into a 96-well plate and were cultured with the indicated compounds for 3 more days (total 12-days of treatment) before cell lysis and quantification of proliferation using the Quant-iT™ PicoGreen dsDNA Assay Kit (ThermoFisher Scientific, Grand Island, NY, USA).

### ChIP-qPCR

A375 cells were grown in complete DMEM media and were incubated with DMSO or A-485 (5 µM) for 12 days. Cells were passaged when they reached confluency. On the 12^th^ day, cells were utilized for ChIP as described previously. ^61,62^ Briefly, cells were fixed in 1% formaldehyde (final concentration) in PBS and chromatin was extracted. The chromatin was then sheared to approximately 200 bp using the Diagenode Bioruptor Pico (Diagenode, Denville, NJ, USA) using 30 cycles (30 seconds on/30 seconds off) according to manufacturer instructions. 1000 µg of sonicated chromatin was then used per immunoprecipitation reaction and were incubated with 4 µg of H3K27ac antibody (4729, Abcam, Cambridge, MA, USA). Immunoprecipitated complexes were washed, the DNA was eluted, treated with Proteinase K, reverse crosslinked and ChIP DNAs were purified using a PCR Cleanup Kit (Qiagen, Germantown, MD, USA). ChIP DNAs were then analyzed for enrichment of H3K27ac at genomic loci via RT-qPCR as described above. ChIP qPCR results were analyzed using the 1% input method. Primer sequences as follows: *THBS1*-Promoter F: CATTCCGGGAGATCAGCTCG, *THBS1*-Promoter R: AAGCATCCCGAAAAGGGACG, *SFRP1*-Promoter F: AGGTGGCTTGGTGTAGAAGC, *SFRP1*-Promoter R: GCGAGTACGACTACGTGAGC, Gene Desert F: AGATGGGTTCACAGTAAGTGGG, Gene Desert R: TATCAGCCGGGTCCCCATTG. Gene Desert located at chr12:61,497,367-61,498,447.

### Transwell invasion assay

A375 and 1205Lu cells were cultured in 10 cm plates and were treated with DMSO or A-485 (5 µM) for 12 days. Cells were passaged when confluent. On the 12^th^ day of treatment, cells were seeded into the transwell invasion assay and allowed to invade for 6 hours. Briefly, 30 µl of solution containing 50 µg of matrigel (Corning, NY, USA) was used to coat the 24-well transwell insert (Corning, NY, USA). 300,000 A375 and 1205Lu cells were seeded into the top chamber of the insert in serum-free DMEM media containing the indicated drugs. Complete DMEM media containing 20% FBS and the indicated drugs was added to the bottom portion of the chamber. After 6 hours of incubation, the transwell inserts were removed, cells were fixed with 70% ethanol for 10 minutes, washed with PBS, stained with propidium iodide (Invitrogen, ThermoFisher Scientific, Carlsbad, CA, USA) and mounted on microscope slides with UltraCruz® mounting media (Santa Cruz Biotechnology, Dallas, TX, USA). 8-10 images were taken on a Nikon E400 (Japan) microscope in a grid pattern of the invaded cells. The number of invaded cells was quantified using ImageJ.

### Statistical analysis

The student’s *t*-test was used for experiments with two experimental groups. ANOVA was used for experiments with more than two experimental groups and was supplemented by the student’s *t*-test as a post hoc test. A *p* value of <0.05 was considered statistically significant. *, **, and *** denote *p* < 0.05, 0.005 and 0.0005, respectively. Standard error of the mean (SEM) was utilized for error bars. Statistical analysis for all experiments were performed on three biological replicates (n=3), unless otherwise indicated. If an alternative statistical test was used for an experiment, it is denoted in the methods section for that experiment.

### Data and code availability

All the RNA sequencing data generated are deposited in the GEO database (accession number GSE254326). Any additional information required can be dire^i^cted to the lead contact. No codes were generated in the manuscript.

## Results

### *EP300* and *SOX10* are commonly co-amplified in human melanomas

Recent studies have found that *EP300* is amplified in acral melanomas at a high frequency. ^32,33^ Intriguingly, *SOX10* is located on 22q13.1, approximately 3 Mb from the *EP300* locus at q13.2 (Figure 1A). Given the proximity of the *EP300* and *SOX10* loci and their significant roles in melanoma development and progression, we sought to explore a functional connection between these two genes in melanoma. Remarkably, we find that *EP300* and *SOX10* gene copy numbers are strongly correlated in human melanomas and are reproducibly co-amplified at a high frequency (∼35% to 46% of samples) across several melanoma datasets from cBioPortal^35,36^, including the TCGA database (Figure 1B). Of note, the high frequency of *EP300/SOX10* co-amplification occurs in both UV-associated and acral tumors (Figure 1B). Copy number gains of *EP300* and *SOX10* were found to correlate with higher gene expression levels, suggesting that these copy number gains are functionally and biologically relevant in tumor samples (Figure 1C). We also found that *EP300* and *SOX10* copy numbers are strongly correlated in human melanoma cell lines using the Cancer Dependency Map^37^ (Figure 1D), suggesting a functional connection between these genes in cellular growth. Furthermore, higher gene copy numbers are associated with higher gene expression of *EP300* and *SOX10* in this setting (Figure 1E). 67.7% (32/51) of melanoma cell lines in the Cancer Dependency Map dataset possessed greater than 2 copies of both *EP300* and *SOX10* and we labeled these lines as EP300/SOX10 coamplified (Figure 1F). These findings were validated using the same pipeline to confirm known co-amplification of *PAK1* and *GAB2* on Chromosome 11 at q14.1 (∼1 Mb apart) in melanoma (Figure S1). ^32^ Importantly, *PAK1/GAB2* and *EP300/SOX10* do not show copy number correlation in either tumors or cell lines using this analysis strategy (Figure S1B-C).

**Figure 1:**
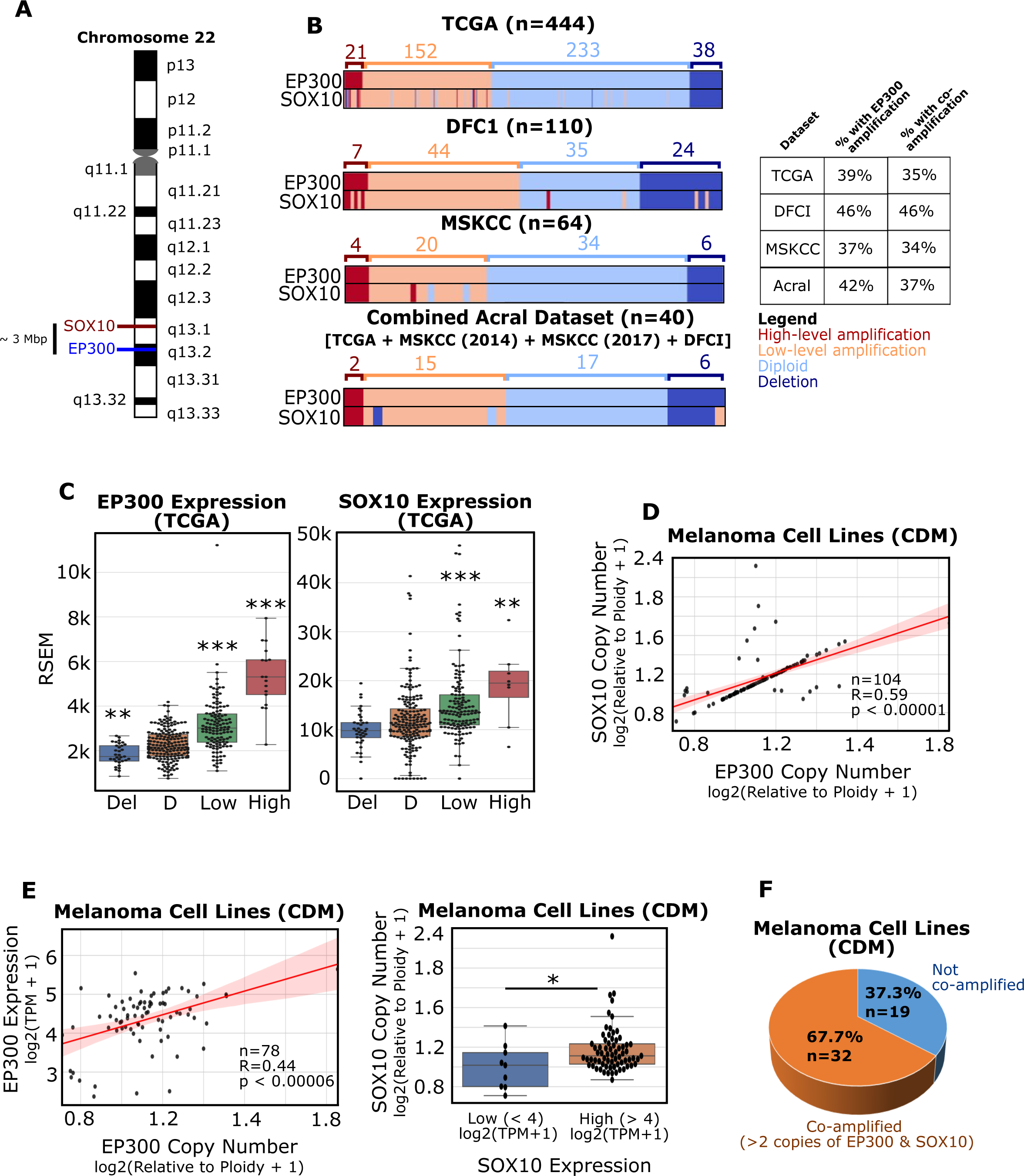
*EP300* is commonly co-amplified with *SOX10* in melanomas. **(A)** The *EP300* and *SOX10* genes are located in close proximity on Chromosome 22. **(B)** *EP300* and *SOX10* amplification prevalence in different melanoma datasets (TCGA, DFCI, MSKCC). Acral patients from multiple datasets were combined (Combined Acral). Amplification levels were defined by GISTIC2.0. **(C)** *EP300* (left) and *SOX10* (right) expression were analyzed for each copy number level, as defined by GISTIC2.0 (data from TCGA; Del: deletion; D: diploid; Low: low-level amplification; High: high-level amplification). **(D)** *EP300* and *SOX10* copy number are positively correlated in melanoma cell lines (data from Cancer Dependency Map [CDM]). **(E)** *EP300* copy number and expression are positively correlated in melanoma cell lines (left). *SOX10* copy number and expression are positively correlated in melanoma cell lines (right) (data CDM). **(F)** *EP300/SOX10* co-amplifications (> 2 copies of *EP300* and *SOX10*) are common in melanoma cell lines (data from CDM). *p < 0.05; **p < 0.005; ***p < 0.0005.

Previous studies have identified higher frequencies of gene amplifications in acral versus UV-associated melanomas^38^; however, this does not appear to be the case for *EP300* (Figure 1B). Rather, the frequency of *EP300* amplification was found to vary greatly between acral melanoma datasets, with 0% of acral samples in the TGEN (2017) dataset possessing *EP300* amplification, despite documented amplification/deletion events in *TERT*, *CCND1*, *CDKN2A* and *NF1* (Figure S2A) ^32^ and higher than 30% *EP300* amplification in other acral datasets (Figure S2B). ^32,33^ Regardless of the frequency of *EP300* gene amplification in acral melanomas, the copy number correlation between *EP300/SOX10* was strongly reproduced across all datasets where *EP300* and *SOX10* were found to be frequently co-amplified (Figure S2C).

### The p300 KAT inhibitor, A-485, downregulates expression of SOX10 protein and associated target genes in human melanoma cells

The high frequency of *EP300/SOX10* co-amplification identified in human melanomas, and previous data from our group demonstrating that EP300 knockdown leads to decreased SOX10 protein levels, ^26^ strongly suggest a functional relationship between *EP300/SOX10*. To investigate this, we classified melanoma cell lines as either *EP300/SOX10* co-amplified (>2 copies of both *EP300/SOX10*) or not co-amplified, treated them with the potent and specific p300 KAT inhibitor, A-485, and probed for SOX10 and downstream SOX10 target genes, such as *MITF* and *DCT*. ^39^ In all cell lines tested, the presence of A-485 led to sharply lower SOX10 protein expression regardless of *EP300/SOX10* amplification status, phenotype (MITF^high^ vs MITF^low^), or melanoma subtype (Figure 2A). In addition, A-485 inhibited MITF and DCT protein and transcript expression level in MITF-high cells (Figures 2A and 2B). In MITF^low^/DCT^low^ A375, Sk-Mel-24 and WM793 melanoma cells without *EP300/SOX10* co-amplification, SOX10 protein expression was also found to be downregulated following treatment with A-485 (Figure 2A). Mining of a SOX10 knockdown dataset^8^ identified *MITF*, *SCD* and *MYC* as being downstream SOX10 effector genes in this melanoma phenotype (Figure 2C). A-485 was found to inhibit *MITF*, *SCD* and *MYC* gene expression in these cell lines (Figure 2D). These results collectively indicate that A-485 potently downregulates SOX10 signaling in human melanoma cells regardless of co-amplification status, melanoma phenotype, or melanoma subtype (UV-associated vs. acral).

**Figure 2:**
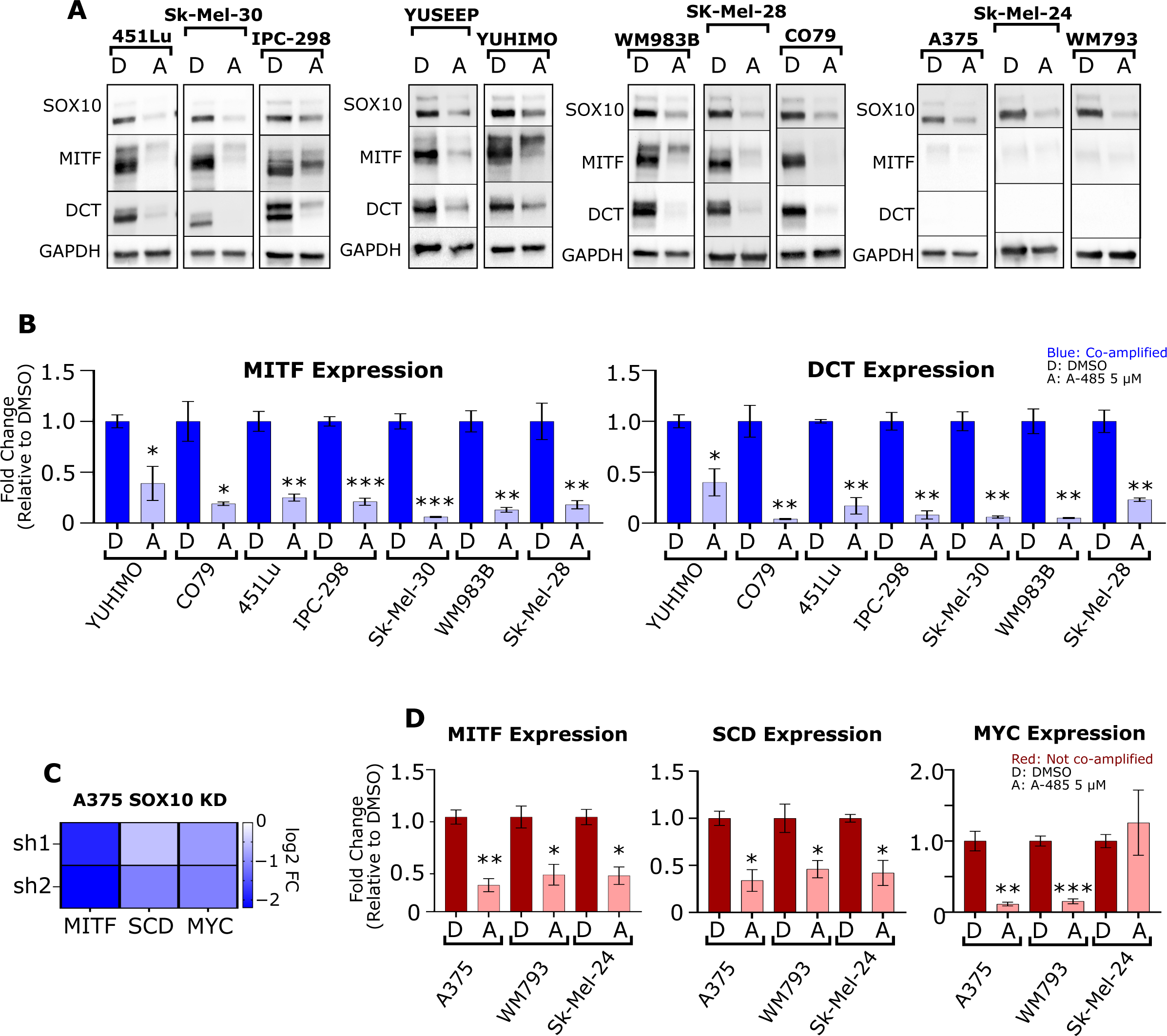
SOX10 signaling requires p300 KAT activity. **(A)** A panel of melanoma cell lines were treated with DMSO (D) or 5 µM A-485 (A) for 24 hours. SOX10, MITF and DCT protein levels were assessed via immunoblotting. **(B)** A panel of *EP300/SOX10* co-amplified melanoma cell lines were treated with DMSO (D) or 5 µM A-485 (A) for 24 hours. *MITF* and *DCT* expression were assessed via RT-qPCR. **(C)** *MITF*, *SCD* and *MYC* expression are downregulated by SOX10 knockdown (KD) in A375 cells. **(D)** A panel of melanoma cell lines without *EP300/SOX10* co-amplifications were treated with DMSO (D) or 5 µM A-485 (A) for 24 hours. *MITF*, *SCD* and *MYC* expression were assessed via RT-qPCR. *p < 0.05; **p < 0.005; ***p < 0.0005.

As we observed a strong association between MITF^high^ expression and *EP300/SOX10* co-amplification status (Figure 2), we sought to determine whether this trend was present in a larger set of melanoma cell lines using the Cancer Dependency Map. ^37^ Indeed, we found that cell lines in the Cancer Dependency Map with *EP300/SOX10* co-amplification also had higher *MITF* expression and lower *AXL* expression than cell lines without *EP300/SOX10* co-amplification (Figure S3A). These results suggest that 22q13.1 and 22q13.2 may contain multiple regulators of the melanocytic, MITF^high^ phenotype. We therefore explored the overlap of genes from these two regions with genes associated with the melanocytic phenotype^40^ and found 18 common genes (Figure S3B). The copy numbers of these 18 genes were statistically correlated in the Cancer Dependency Map (Figure S3C), suggesting that this entire region may be linked. Furthermore, many of these genes showed a statistically significant correlation between copy number and expression in the Cancer Dependency Map (Figure S3D). In particular, SREBF2 controls the expression of SCD, a known regulator of MITF^41,42^, suggesting that 22q13.1 and 22q13.2 cytobands contain a cluster of melanocytic phenotype genes with known roles in transcriptional activation of MITF including p300, SOX10, and SREBF2 (Figure S3E).

### A-485 induces proteasomal degradation of SOX10 in human melanoma cells

As p300 KAT inhibition with A-485 led to reduced expression of SOX10 target gene transcripts and SOX10 protein (Figure 2) we next sought to determine the effects of A-485 on *SOX10* transcript levels. Remarkably, we found that A-485 does not significantly downregulate *SOX10* mRNA levels in 9 of 10 melanoma cell lines tested (Figure 3A), which was confirmed using alternative *SOX10* primers (Figure S4A). Downregulation of SOX10 protein was confirmed with an alternative SOX10 antibody (Figure S4B). Query of the p300DB regulome database^28^ found that *p300* knockout and A-485 treatment reduced SOX10 protein levels, but not *SOX10* mRNA expression in MEF cells (Figure S4C), further confirming our results. The p300 bromodomain inhibitor CCS1477 also reduced SOX10 protein levels in A375 and IPC-298 cells (Figure S4D). Importantly, A-485 and CCS1477 are structurally unique compounds that target distinct domains on p300, suggesting that downregulation of SOX10 protein expression is a specific on-target effect of p300 inhibition which downregulates SOX10 protein expression through a post-transcriptional mechanism.

**Figure 3:**
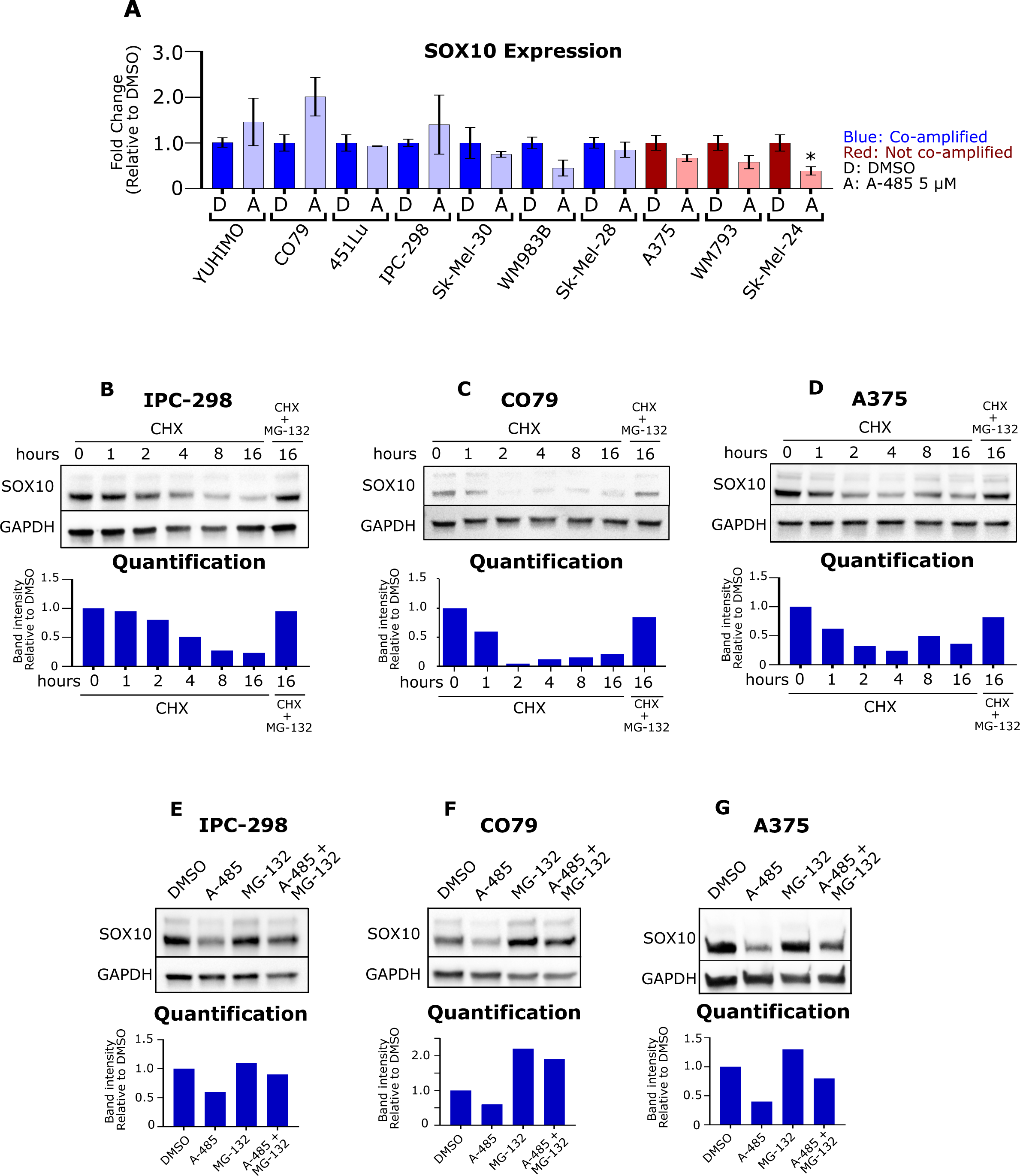
A-485 induces proteasomal degradation of SOX10. **(A)** A panel of melanoma cell lines were treated with DMSO (D) or 5 µM A-485 (A) for 24 hours. *SOX10* expression was assessed via RT-qPCR. **(B-D)** IPC-298, CO79 and A375 cells were treated with 50 µg/mL cyclohexamide and/or 5 µM MG-132 for the indicated time. SOX10 protein levels were assessed via immunoblotting. **(E-G)** IPC-298, CO79 and A375 cells were treated with DMSO, 5 µM A-485, 5 µM MG-132, or their combination for 16 hours. SOX10 protein levels were assessed via immunoblotting. *p < 0.05.

To explore this hypothesis, we treated IPC-298, CO79 and A375 cells with cycloheximide (CHX) and/or the proteasome inhibitor, MG-132, to evaluate SOX10 protein stability and targeted degradation by the proteasome(Figure 3B-D). ^20^ Notably, we find that SOX10 is degraded over time following treatment with CHX and that this degradation can be rescued by MG-132 (Figure 3B-D). We next evaluated whether MG-132 could restore SOX10 protein levels in the setting of A-485. Remarkably, we found that MG-132 was able to rescue A-485 effects on SOX10 protein levels in all three cell lines (Figure 3E-G), suggesting that inhibition of p300 KAT activity promotes proteasomal degradation of SOX10.

### A-485 potently and preferentially inhibits expression of SOX10 target genes in human melanoma

As A-485 was found to induce degradation of SOX10 protein levels without associated changes in *SOX10* transcription, we sought to determine the broader transcriptional consequences of A-485 on melanoma cells and to assess the expression of the global SOX10 gene network. IPC-298, CO79 and A375 cells were treated with 5 uM A-485 for 24 hours and evaluated by RNA-seq. As anticipated, A-485 led to downregulated expression of a large number of genes given its function as a transcriptional coactivator (Figure 4A-C). The top five Gene Ontology (GO) ^43^ pathways for genes downregulated by A-485 in all three cell lines included changes in pathways associated with cellular differentiation, cell adhesion, and cell motility (Figure 4D).

**Figure 4:**
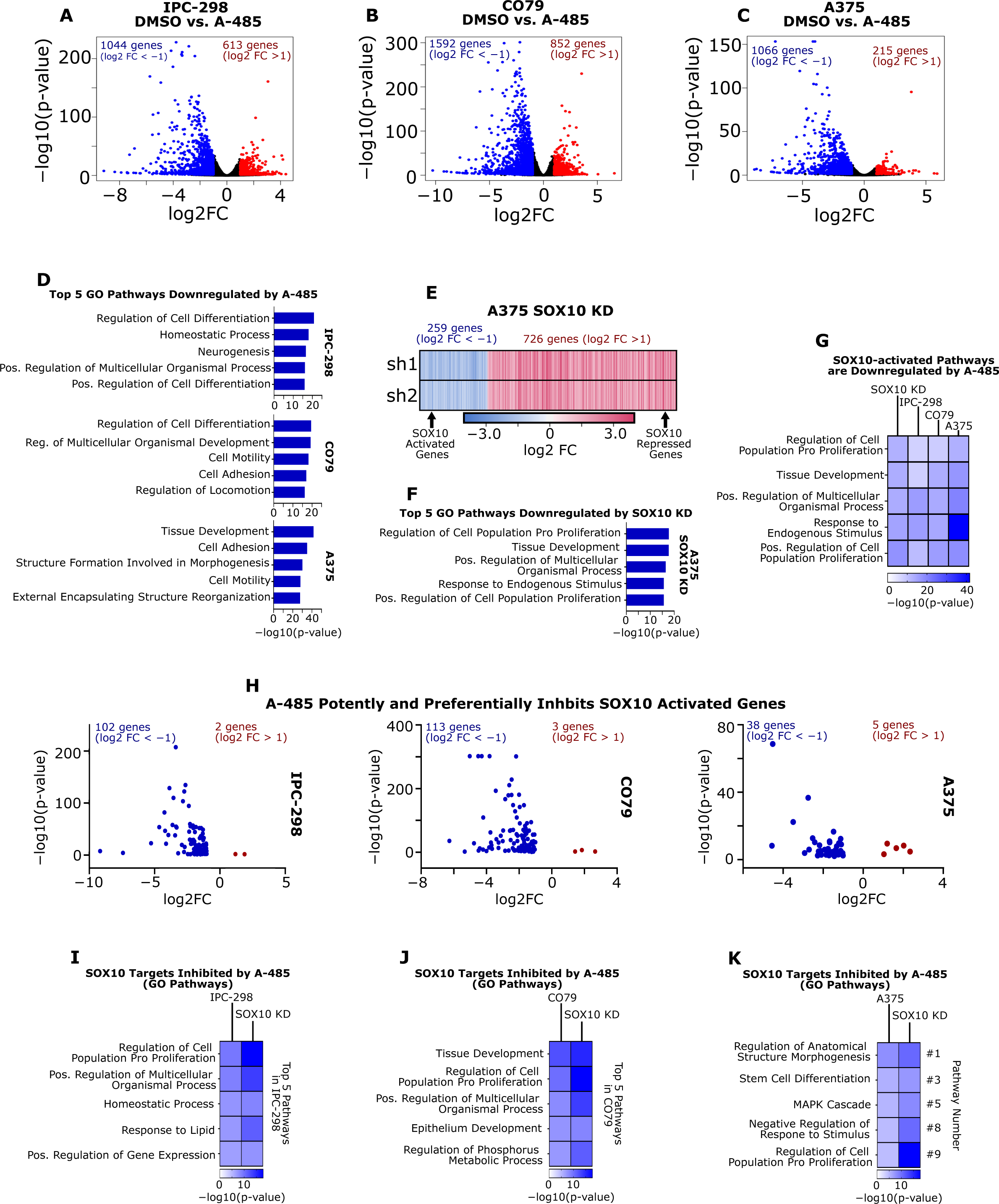
A-485 treatment potently downregulates SOX10 target genes in melanoma cells. **(A-C)** IPC-298, CO79 and A375 cells were treated with 5 µM A-485 for 24 hours and RNA-sequencing was performed. Volcano plots of differentially expressed genes in each cell line are depicted (DEG defined as FC > |2| and p < 0.05). **(D)** The top five Gene Ontology Biological Process pathways are shown for downregulated genes in each cell line. **(E)** A heatmap of DEGs resulting from SOX10 KD in A375 cells is shown (data from GSE50535). **(F)** The top five Gene Ontology Biological Process pathways are shown for SOX10-activated genes (i.e. genes downregulated by SOX10 KD). **(G)** A-485 downregulates Gene Ontology Biological Process pathways that are activated by SOX10 in IPC-298, CO79 and A375 cells. **(H)** Volcano plots of SOX10-activated genes differentially expressed due to A-485 treatment for IPC-298, CO79 and A375 cells. **(I-K)** Gene Ontology Biological Process pathways are shown for SOX10-activated genes that are downregulated by A-485.

We next sought to determine the impact of A-485 treatment of melanoma cells on SOX10 target genes using the SOX10 KD dataset in A375 cells^8^ and classified the differentially expressed genes as either SOX10-activated (genes downregulated by KD) or SOX10-repressed (genes activated by KD) (Figure 4E). The top five GO pathways for SOX10-activated genes include pathways related to proliferation and development (Figure 4F), which are GO pathways that are downregulated by A-485 treatment in IPC-298, CO79 and A375 cells (Figure 4G). This suggests that SOX10 is a major target of p300 KAT inhibition in human melanoma (Figures 4G). Of note, A-485 downregulated a large subset of SOX-10 activated genes in IPC-298, CO79 and A375 cells (Figure 4H), and these A-485-repressed, SOX-10-activated genes were enriched for pathways involved in cell proliferation and differentiation (Figures 4I-K). These data were validated by an independent dataset of Sk-Mel-5 cells treated with A-485 (GSE116459) and our previously published EP300 KD datasets in WM983B and SK-Mel-5 cells (Figure S5A-C). ^26,34^ with strong congruence between RNA-seq and qPCR results for SOX10 target genes (Figures S5D, S5E).

### Inhibition of p300 KAT activity leads to growth inhibition in SOX10+ melanoma cells

Our lab and others have reported growth inhibition of melanoma cell lines in response to A-485 treatment in an MITF-dependent manner. ^26,44^ However, MITF^low^ cells can be SOX10+ and require SOX10 for growth. ^8,13^ We hypothesized that SOX10+/MITF^low^ cells could be responsive to A-485, but would require a longer treatment period to impact downstream SOX10 effectors beyond MITF. To investigate this, we performed 6-day A-485 dose-response assays on a panel of melanoma cell lines with cells classified as *EP300/SOX10* co-amplified/MITF^high^ and *EP300/SOX10* non-amplified/MITF^low^ cells (Figure 5A) and found that co-amplified/MITF^high^ cells were potently growth inhibited by A-485, relative to the non-amplified/MITF-low cells (Figure 5A), including the acral melanoma cell lines CO79, YUHIMO and YUSEEP. Next, we investigated whether melanoma cells that did not respond to 6-day treatment with A-485 (short-term assay) would be growth inhibited by 12-days of treatment (long-term assay). Remarkably, Sk-Mel-28, A375, WM793 and Sk-Mel-24 cells demonstrated significant growth inhibition following long-term exposure to A-485 (Figure 5B) suggesting that chromatin remodeling by p300 KAT activity may involve downstream effector pathways in SOX10+/MITF^low^ cells.

**Figure 5:**
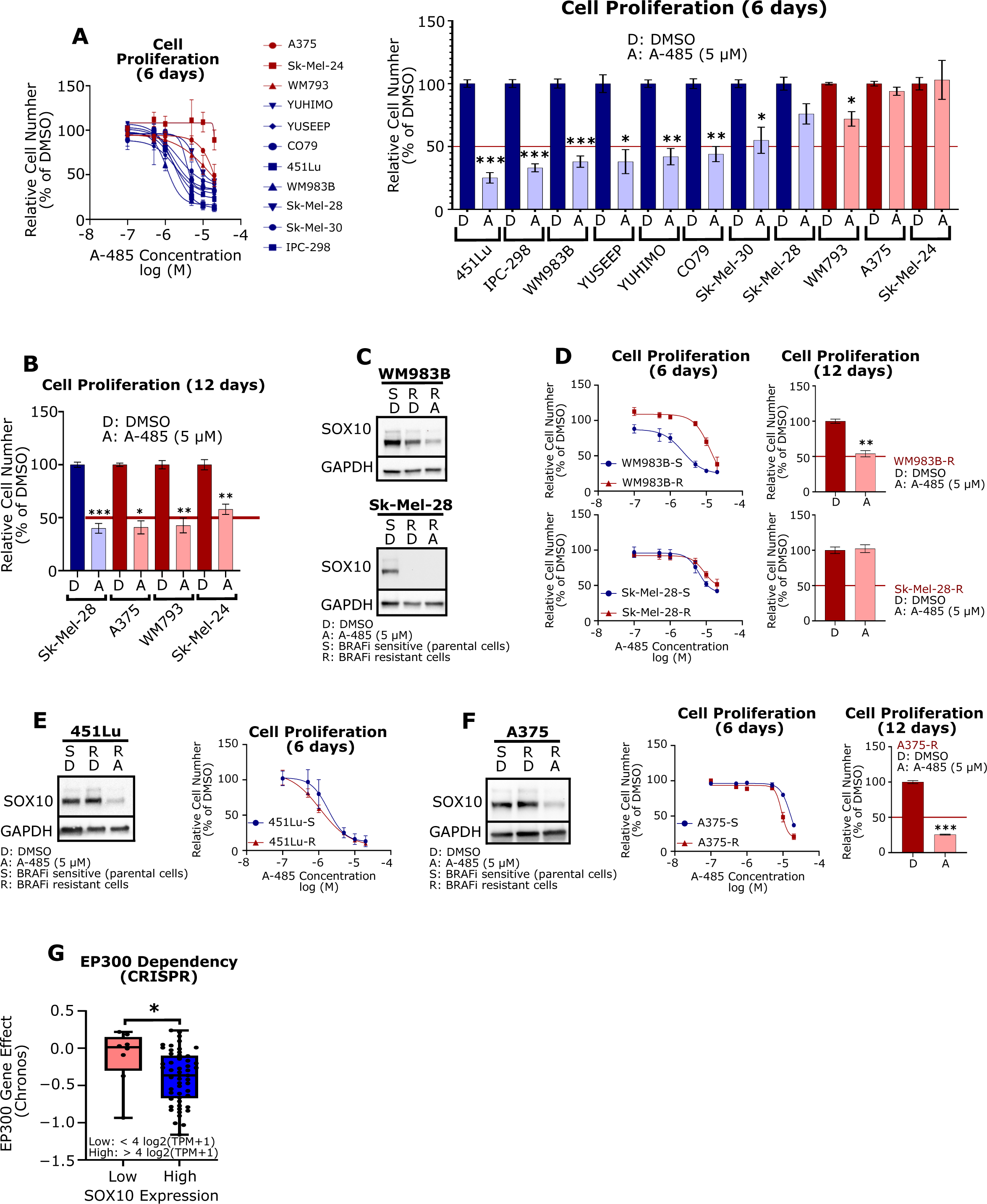
A-485 potently inhibits proliferation of SOX10+ melanoma cell lines. **(A)** A panel of melanoma cell lines were treated with A-485 (in a range of doses) for 6 days and their relative proliferation versus control was assessed (left). Results for 5 µM A-485 versus control are shown (right). **(B)** Cells that did not respond to A-485 within 6 days were pre-treated with 5 µM A-485 for 9 days and re-seeded for a 3-day proliferation assay and their relative proliferation versus control was assessed (12 days total of drug treatment). **(C)** WM983B-R and SK-Mel-28-R BRAF inhibitor resistant cells have low expression of SOX10 in comparison to parental cells. **(D)** The proliferation effects of short-term (6-day) or long-term (12-day) A-485 treatment of WM983B-R and SK-Mel-28-R cells was assessed. **(E)** 451Lu-R cells maintain SOX10 expression versus BRAFi-sensitive parental cells (left). 451Lu-S and 451Lu-R were treated with A-485 (in a range of doses) for 6 days and their relative proliferation versus control was assessed (right). **(F)** A375-R cells maintain SOX10 expression versus BRAFi-sensitive parental cells (left). A375-S and A375-R were treated with A-485 (in a range of doses) for 6 days and their relative proliferation versus control was assessed (middle). A375-S and A375-R cells were pre-treated 5 µM A-485 for 9 days and re-seeded for a 3-day proliferation assay and their relative proliferation versus control was assessed (right). **(G)** EP300 dependency scores are plotted for melanoma cell lines with low SOX10 (<4 log2(TPM+1) versus high SOX10 (>4 log2(TPM+1) expression in the CDM 22Q2 Public+Score, Chronos dataset. *p < 0.05; **p < 0.005; ***p < 0.0005.

Loss of SOX10 expression is a well-characterized mechanism of resistance to MAPKi. ^13^ We therefore explored A-485 effects on the MAPKi-resistant cell lines WM983B BRAFi-R (partial loss of SOX10) and SK-Mel-28 BRAFi-R cells (SOX10 undetectable) (Figure 5C) to determine whether A-485 could effectively inhibit tumor cell growth in this setting. Not surprisingly, WM983B BRAFi-R (SOX10-low) cells were less responsive to A-485 in the short term versus their BRAFi-S parental cells (SOX10-high) (Figure 5D); however, WM983B BRAFi-R (SOX10-low) cells eventually responded to A-485 following long-term exposure (Figure 5D). Sk-Mel-28 BRAFi-R cells (SOX10 undetectable) were found to be highly resistant to A-485 effects on tumor cell growth following both short-term (6-day) and long-term (12-day) time points (Figure 5D); however, the BRAFi-sensitive parental Sk-Mel-28 BRAFi-S (SOX10+) cells were sensitive to A-485 growth inhibition following long-term (12-day) treatment (Figure 5B), suggesting that complete loss of SOX10 in Sk-Mel-28 BRAFi-R cells rendered them insensitive to A-485.

In order to explore whether changes in SOX10 expression drive resistance to A-485 versus BRAFi-resistance alone, we explored the effect of A-485 in 451Lu BRAFi-R and A375 BRAFi-R cells which maintain equivalent SOX10 expression levels with their parental BRAFi-S cells. Remarkably, we found no differential response to A-485 in these BRAFi-S and BRAFi-R cells (Figures 5E, 5F). These results suggest that A-485 specifically inhibits SOX10-dependent proliferation in melanoma regardless of BRAFi sensitivity. A search of the Cancer Dependency Map further confirmed that p300 drives proliferation in a SOX10-dependent manner as melanoma cells with high SOX10 expression are more dependent on p300 than those with low SOX10 expression (Figure 5G).

### A-485 potently inhibits invasion in MITF^low^ melanoma cells

While we have clearly demonstrated that p300 inhibition downregulates expression of SOX10-activated genes and SOX10-dependent proliferation in human melanoma cells, it is important to recognize that SOX10 loss-of-function is also correlated with increased melanoma invasive capacity through upregulation of a metastatic transcriptional program that defines the undifferentiated phenotype. ^13^ We hypothesized that a subset of SOX10-repressed genes (i.e. genes activated upon SOX10 KD) would require p300 for their activation given the critical role of p300 as a broadly effective transcriptional coactivator (Figure 6A). ^27^ Remarkably, we found that a subset of SOX10-repressed genes were also repressed by A-485 treatment in IPC-298, CO79 and A375 cells (Figure 6B). Notably, we determined that the SOX10-repressed genes which are also repressed by A-485 are strongly enriched for GSEA^45,46^ and KEGG invasion-related pathways in A375 cells (Figure 6C). A-485 also downregulates the expression of many genes that are part of the overall GSEA EMT Hallmark gene set (Figure 6D), including genes that are downstream targets of SOX10 (Figure 6E). Similar results were seen in MITF^high^ IPC-298 and CO79 cells (Figure S6A). These results were confirmed by RT-qPCR in A375, WM793, Sk-Mel-24 and 1205Lu cells (Figure S6B-D). A-485 was also found to potently and preferentially inhibit expression of genes involved in KEGG invasion-related pathways, such as ECM Interaction, Focal Adhesion, Regulation of Actin Cytoskeleton^47,48^, Cell Adhesion Molecules and Axon Guidance^49^ (Figure 6F) as well as expression of 13 collagen genes associated with melanoma invasion (Figure 6F). ^50^

**Figure 6:**
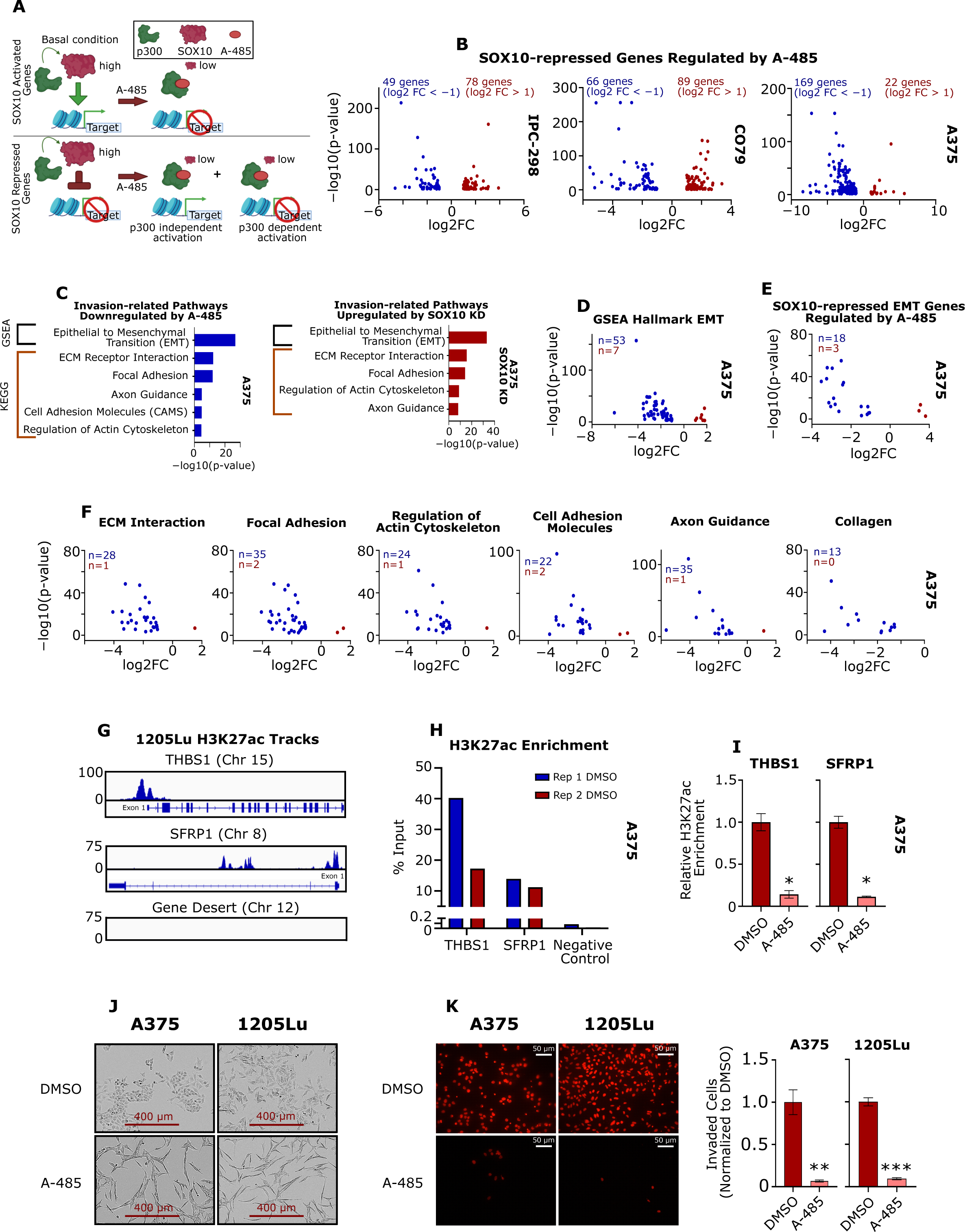
p300 KAT activity is essential for activation of SOX10-repressed EMT markers. **(A)** Diagram depicting the hypothesis that p300 activity is essential for expression of SOX10-activated genes, but SOX10-repressed genes may also require p300 for their activation. **(B)** Volcano plots of SOX10-repressed genes differentially expressed due to A-485 treatment for IPC-298, CO79 and A375 cells (DEG defined as FC > |2| and p < 0.05). **(C)** SOX10-repressed genes are enriched for genes involved in the epithelial-to-mesenchymal transition (EMT) and similar invasion-related pathways are downregulated by A-485. **(D)** Volcano plot of EMT genes differentially expressed due to A-485 treatment for A375 cells. **(E)** Volcano plot of SOX10-regulated EMT genes differentially expressed due to A-485 treatment for A375 cells. **(F)** Volcano plots of genes differentially expressed due to A-485 are shown for various KEGG invasion-related pathways identified in (C). Differentially expressed collagen genes are also shown. **(G)** H3K27ac is enriched at the promoters of EMT genes THBS1 and SFRP1, in comparison to a gene desert in invasive 1205Lu cells via ChIP-seq. **(H)** H3K27ac was confirmed to be enriched at the promoters of *THBS1* and *SFRP1*, in comparison to a gene desert in A375 cells via ChIP-qPCR. **(I)** A-485 decreases H3K27ac enrichment at the promoters of *THBS1* and *SFRP1* as assessed by ChIP-qPCR. **(J)** 5 µM A-485 alters the cell morphology of A375 and 1205Lu cells after long-term (12-day) treatment. (K) A-485 potently inhibits invasion in A375 and 1205 after long-term (12-day) treatment. Representative images of invaded cells are shown on the left and quantification of invaded cells is shown on the right. *p < 0.05; **p < 0.005; ***p < 0.0005.

In order to confirm p300 regulation of EMT-associated genes independent of SOX10, we explored p300-associated H3K27 histone acetylation at EMT-associated gene regulatory regions. ^28,51–53^ H3K27ac levels at active regulatory regions are known to be sensitive to A-485 inhibition. ^28,44,52^ H3K27ac ChIP-seq was previously performed in 1205Lu-R cells (GSE254703) and we selected EMT genes enriched for H3K27ac which were also downregulated by A-485 in our RNA-seq and qPCR data for further investigation. For example, *THBS1* and *SFRP1* were highly enriched for H3K27ac at their promoters versus a gene desert on Chromosome 12 in 1205Lu-R cells (Figure 6G) and expression of these genes was downregulated by A-485 (Figure S6B-D). We confirmed H3K27ac enrichment at the *THBS1* and *SFRP1* promoters in A375 cells by ChIP-qPCR (Figure 6H) and A-485 was found to strongly decrease H3K27ac enrichment at these promoter sites (Figure 6I). These results suggest that p300 acetylation of chromatin at invasion-related genes is required for activating these genes upon SOX10 loss-of-function.

Finally, we noticed that A-485 induces a significant change in cellular morphology of A375 and 1205Lu cells during long-term (12-day) treatment (Figures 6J, S6E). We hypothesized that this change in cellular morphology may correlate with decreased invasive potential. Indeed, we found that A-485 significantly decreased cellular invasion of A375 and 1205Lu cells in a Matrigel® transwell assay (Figure 6K).

## Discussion

SOX10 is a neural crest lineage-specific transcription factor that regulates melanoma development, rapid tumor growth and tumor immunogenicity. ^4–12^ Additionally, SOX10 has been shown to play a crucial role in melanoma phenotype switching through repression of a metastatic transcriptional program. ^13^ Efforts to therapeutically target SOX10 have been challenging due to the relative inaccessibility of targeted therapies to transcription factors and specific complexities associated with loss of SOX10 activity, which may promote tumor cell invasion and metastasis.

As a transcription factor, SOX10 relies on chromatin remodelers and epigenetic enzymes to regulate both SOX10-activated and SOX10-repressed target genes^54–57^; thus, key epigenetic regulators of SOX10 signaling potentially represent a novel therapeutic strategy to target SOX10 function. In this study, we show that the transcriptional coactivator and epigenetic lysine acetyltransferase p300 is critical for promoting SOX10 protein stability and that chemical inhibition of p300 KAT activity by A-485 results in potent downregulation of SOX10 protein levels through proteasomal degradation, and expression of SOX10-repressed genes.

Downregulation of SOX10-activated genes by A-485 leads to inhibition of melanoma cell growth as well as downregulated expression of invasion-related genes in MITF^low^ melanoma cells, including EMT genes that are repressed by SOX10.

Our results collectively demonstrate that A-485 represents an attractive therapeutic tool to target both SOX10-dependent proliferation and SOX10-independent invasion with a single agent. Furthermore, our results delineate p300 as a novel and critical activator of the metastatic transcriptional program that defines MITF-low cells.

The duality of p300 regulating proliferation in MITF-high cells, but strongly promoting invasion in MITF-low cells is intriguing. We hypothesize that, like other models, p300 binding is redistributed when a prominent transcription factor (such as SOX10 or MITF) is either activated or inhibited. ^58^ This redistribution could relocate p300 from proliferation genes to invasion-related genes in MITF-high cells compared to MITF-low/SOX10 negative cells, respectively. This redistribution could also partially explain why MITF-low cells are less-responsive to A-485 in terms of proliferation, but potently respond to A-485 in terms of invasion. In support of this hypothesis, SOX10 is known to bind active areas of chromatin marked by H3K27ac^57,59^, which is specifically catalyzed by p300. ^51^ This strongly indicates p300 and SOX10 share genomic binding sites and that p300 may regulate SOX10 signaling through several mechanisms and that loss of SOX10 or MITF could cause p300 redistribution to alternative epigenetic regulatory regions. Indeed, evidence suggests that many SOX10 binding sites are co-bound by p300, but future work will be needed in human cells to fully explore the functional relationship between p300 and SOX10.

Finally, for p300 inhibitors to be effective in the clinic, it would be useful to have a biomarker for therapeutic sensitivity. Our data show that *EP300* and *SOX10* are commonly co-amplified in both UV-associated and acral melanoma tumors. These amplifications appear to be functionally and biologically relevant because they confer higher expression of both genes. Other groups have proposed *EP300* amplifications may serve as a biomarker for sensitivity to p300 inhibitors in the clinic^33^; however, our results suggest that *EP300/SOX10* co-amplifications could serve as a more comprehensive marker of sensitivity versus EP300 amplifications alone, given the implications for SOX10 degradation and SOX10-independent invasion. Our work also demonstrates that MITF^low^ melanoma cell lines are highly dependent on p300 for expression of invasion-related genes. MITF^low^ melanoma status may therefore serve as a biomarker for tumors which would be most sensitive to inhibition of invasion and metastasis through p300 KAT inhibition. Overall, our results show a novel p300 and SOX10 regulatory axis and suggest that p300 KAT inhibition may confer anti-cancer effects to heterogeneous populations of melanoma cells.

## Acknowledgments

Funding from the NIH R35GM149229 partly supported this work. We thank Dr. Sam Whedon for helpful advice and Drs. Meenard Herlyn, Anurag Singh, Nick Hayward, Ruth Halaban, Deborah Lang, and Jong-In Park for providing cells for these experiments. We thank the Queensland Institute of Medical Research (QIMR Berghofer) and the University of Colorado Skin Cancer Biorepository for providing acral melanoma cell lines for this research.

## Author contributions

R.M.A. provided the overall supervision of the project. A.W., M.C., P.A.C, and R.M.A contributed to conceptualization of different parts of the project and contributed to experiment design. A.W., N.G, K.L., W.A.W., S.C, and K.P. conducted experiments. A.W. performed the data analysis.A.W. wrote the manuscript, and the remaining authors reviewed and edited the manuscript to generate the final version.

