## Supplemental Figures for "p300 KAT regulates SOX10 stability and function in human melanoma"

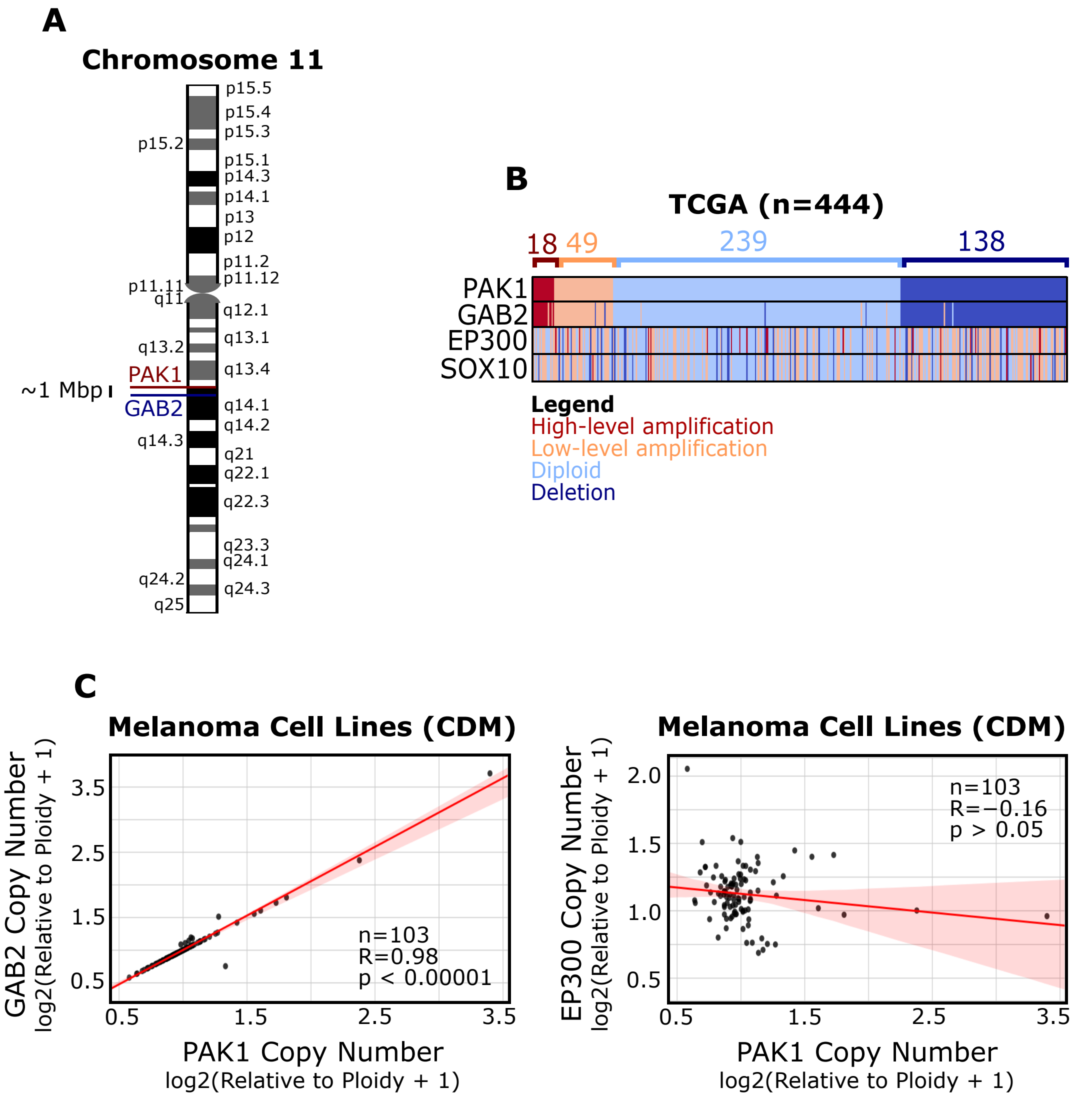
A
Chromosome 11
p15.5
p15.4
p15.3
p15.2
p15.1
p14.3
p14.1
p13
p12
p11.2
p11.12
p11.11
q11
q12.1
q13.1
q13.2
q13.4
PAK1
~1 Mbp
GAB2
q14.1
q14.2
q14.3
q21
q22.1
q22.3
q23.3
q24.1
q24.2
q24.3
q25
B
TCGA (n=444)
18
49
239
138
PAK1
GAB2
EP300
SOX10
Legend
High-level amplification
Low-level amplification
Diploid
Deletion
C
Melanoma Cell Lines (CDM)
GAB2 Copy Number
log2(Relative to Ploidy + 1)
PAK1 Copy Number
log2(Relative to Ploidy + 1)
n=103
R=0.98
p < 0.00001
Melanoma Cell Lines (CDM)
EP300 Copy Number
log2(Relative to Ploidy + 1)
PAK1 Copy Number
log2(Relative to Ploidy + 1)
n=103
R=−0.16
p > 0.05

**Supplementary Figure 1: The copy numbers of PAK1 and GAB2 correlate with each other but not with EP300 or SOX10. (A)** The PAK1 and GAB2 genes are located closely together on Chromosome 11. **(B)** PAK1 and GAB2 copy number levels are correlated in TCGA tumor samples, but they do not correlate with copy number levels of EP300 or SOX10. Copy number levels were defined by GISTIC2.0. **(C)** PAK1 and GAB2 copy numbers are positively correlated in melanoma cell lines (data from Cancer Dependency Map [CDM]). **(D)** EP300 and PAK1 copy numbers are not correlated in melanoma cell lines (data from Cancer Dependency Map [CDM]).

A

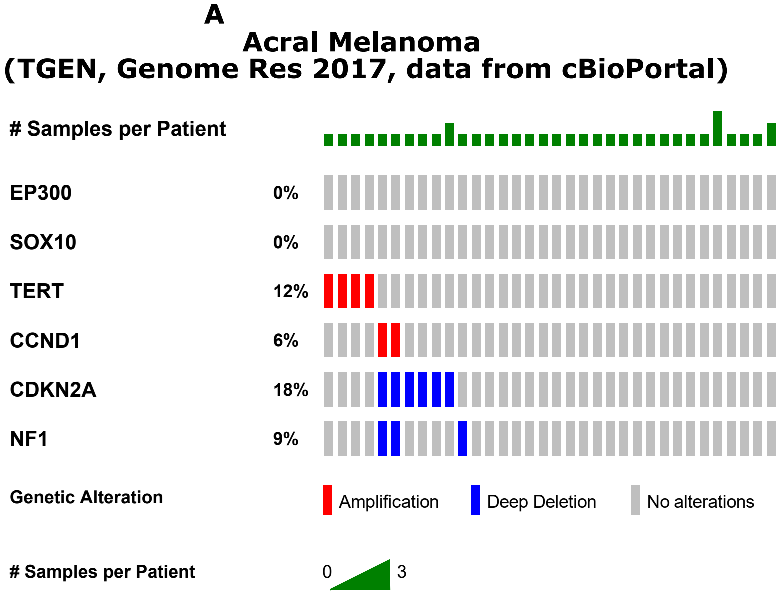


B

| Acral Datasets | % with EP300 amplification |
|---|---|
| TGEN (2017) | 0% 0/38 samples |
| Yeh et al. (2019) | 16.4% 20/122 samples |
| Shi et al. (2022) | 31.7% 19/60 samples |

C

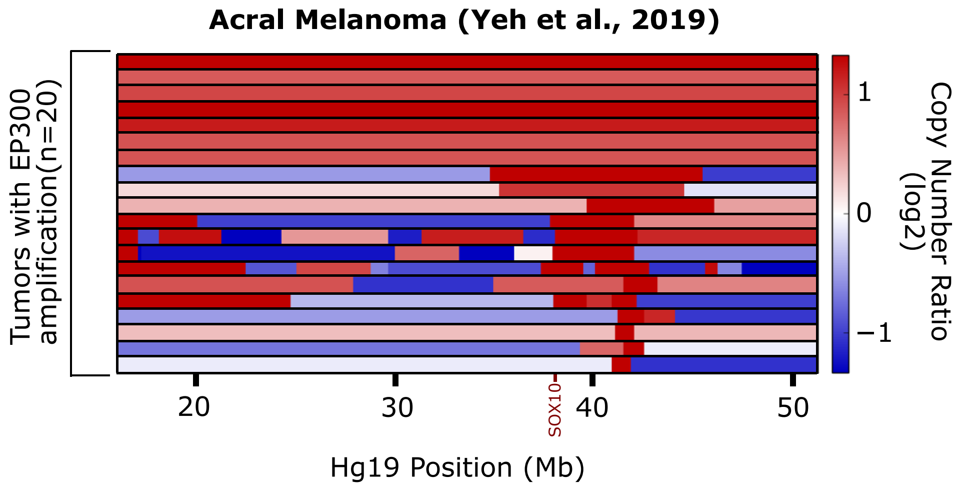

**Supplementary Figure 2: EP300 amplification frequency is widely variable in acral melanoma datasets, but EP300/SOX10 co-amplifications are reproducible. (A)** OncoPrint is shown for the TGEN Acral Melanoma Dataset (2017). Dataset is retrieved from cBioPortal. Copy number levels defined by GISTIC2.0. **(B)** Table of EP300 amplification frequencies in several acral melanoma datasets. **(C)** EP300 and SOX10 copy numbers from acral patients in the Yeh et al. (2019) dataset.

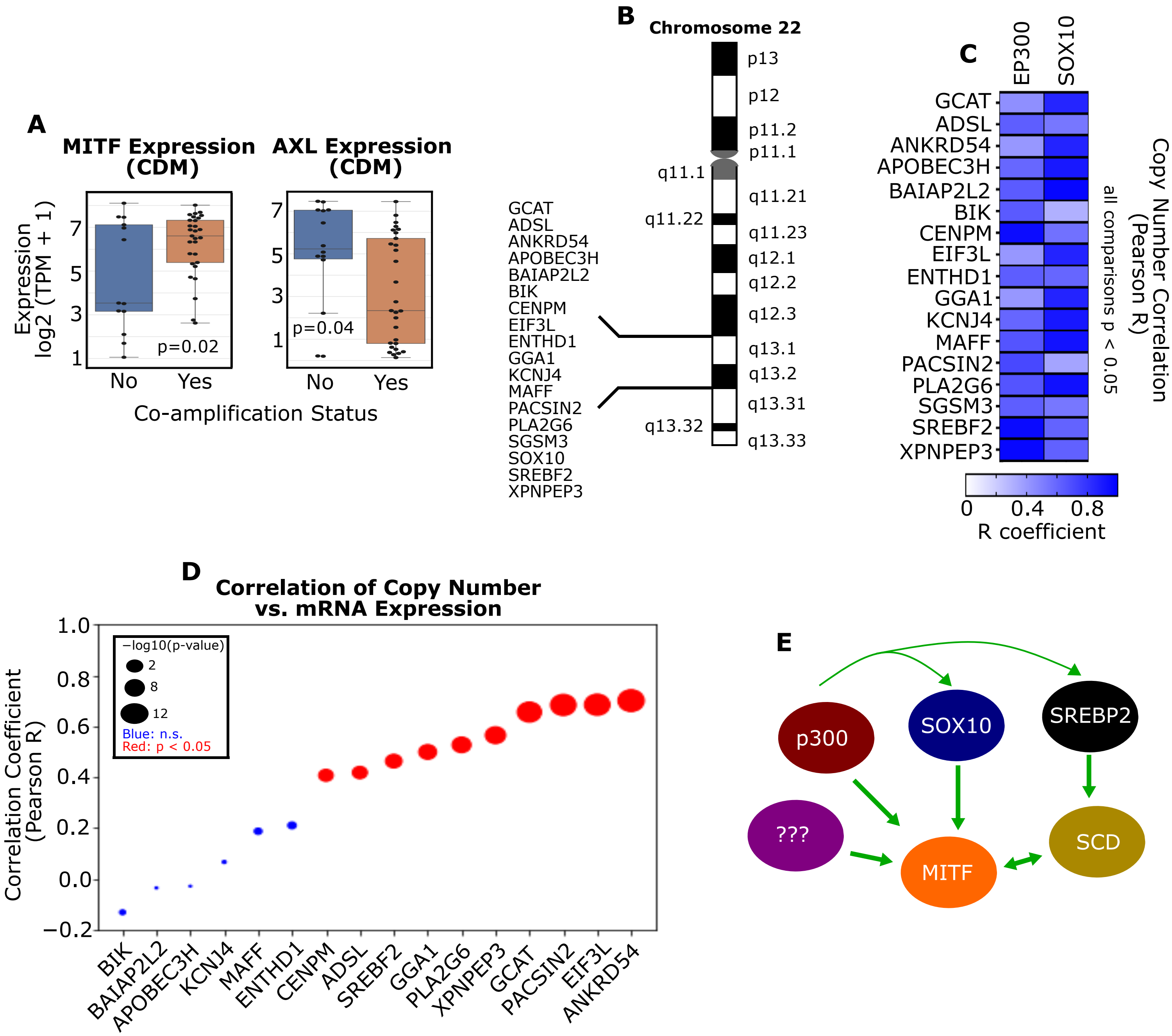
A
MITF Expression (CDM)
AXL Expression (CDM)
Expression log2 (TPM + 1)
p=0.02
p=0.04
No
Yes
Co-amplification Status
B
Chromosome 22
p13
p12
p11.2
p11.1
q11.1
q11.21
q11.22
q11.23
q12.1
q12.2
q12.3
q13.1
q13.2
q13.31
q13.32
q13.33
GCAT
ADSL
ANKRD54
APOBEC3H
BAIAP2L2
BIK
CENPM
EIF3L
ENTHD1
GGA1
KCNJ4
MAFF
PACSIN2
PLA2G6
SGSM3
SOX10
SREBF2
XPNPEP3
C
EP300
SOX10
Copy Number Correlation (Pearson R)
all comparisons p < 0.05
R coefficient
D
Correlation of Copy Number vs. mRNA Expression
Correlation Coefficient (Pearson R)
−log10(p-value)
Blue: n.s.
Red: p < 0.05
E
p300
SOX10
SREBP2
???
MITF
SCD

**Supplementary Figure 3: Chromosome 22 q13.1 and q13.2 regions contain a melanocytic cluster of genes. (A)** Cell lines with EP300/SOX10 co-amplifications have higher MITF and lower AXL expression than cell lines without co-amplifications (data from Cancer Dependency Map [CDM]). **(B)** q13.1 and q13.2 on Chromsome 22 contain 18 genes associated with the melanocytic phenotype in melanoma. **(C)** The copy number of the 18 melanocytic genes in q13.1 and q13.2 are positively correlated (data from Cancer Dependency Map [CDM]). **(D)** The correlation coefficient R is shown for the correlation of copy number versus mRNA expression for the 18 melanocytic genes (data from Cancer Dependency Map [CDM]). **(E)** At least EP300, SOX10 and SREBF2 genes are known regulators of MITF function in melanoma.

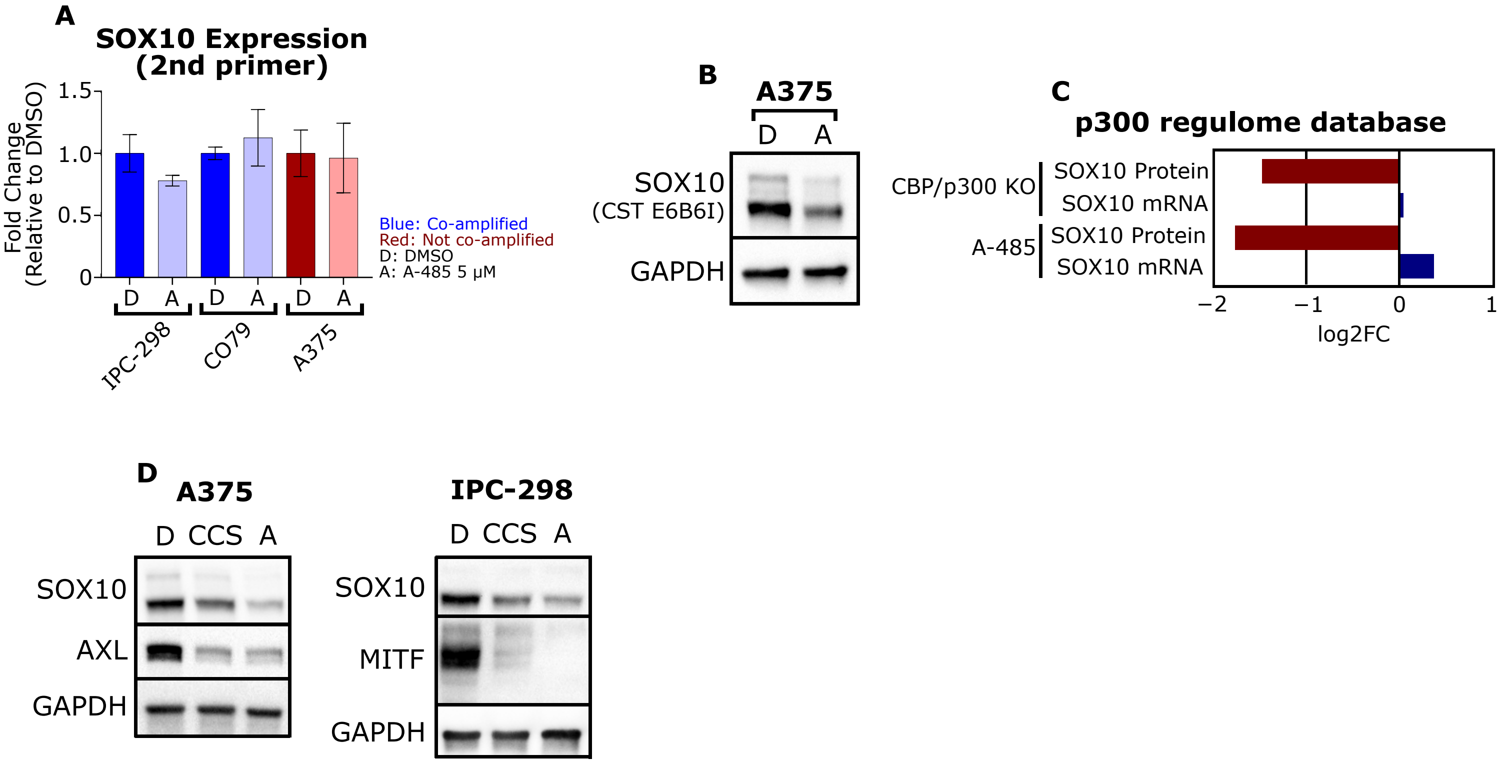
A
SOX10 Expression (2nd primer)
Fold Change (Relative to DMSO)
1.5
1.0
0.5
0
D
A
IPC-298
CO79
A375
Blue: Co-amplified
Red: Not co-amplified
D: DMSO
A: A-485 5 μM
B
A375
SOX10
(CST E6B6I)
GAPDH
C
p300 regulome database
CBP/p300 KO
A-485
SOX10 Protein
SOX10 mRNA
−2
−1
0
1
log2FC
D
A375
IPC-298
D
CCS
A
SOX10
AXL
MITF
GAPDH

**Supplementary Figure 4: SOX10 is not transcriptionally downregulated by A-485, but is decreased at the protein level. (A)** RT-qPCR validation of SOX10 mRNA expression following 5 µM A-485 treatment using a second qPCR primer. Data are represented as mean ± SEM. (B) Immunoblot validation of SOX10 protein levels following A-485 treatment using a second SOX10 antibody (CST 69661). D: DMSO, A: 5 µM A-485 **(C)** A-485 treatment and CBP/p300 knockout (KO) results in downregulation of SOX10 protein levels, but not decreased SOX10 mRNA expression in MEFs (data from p300 regulome database). **(D)** p300 bromodomain inhibitor CCS1477 (5 µM) downregulates SOX10 protein levels in melanoma cells and recapitulates the effects of A-485 on AXL and MITF in A375 and IPC-298 cells, respectively.

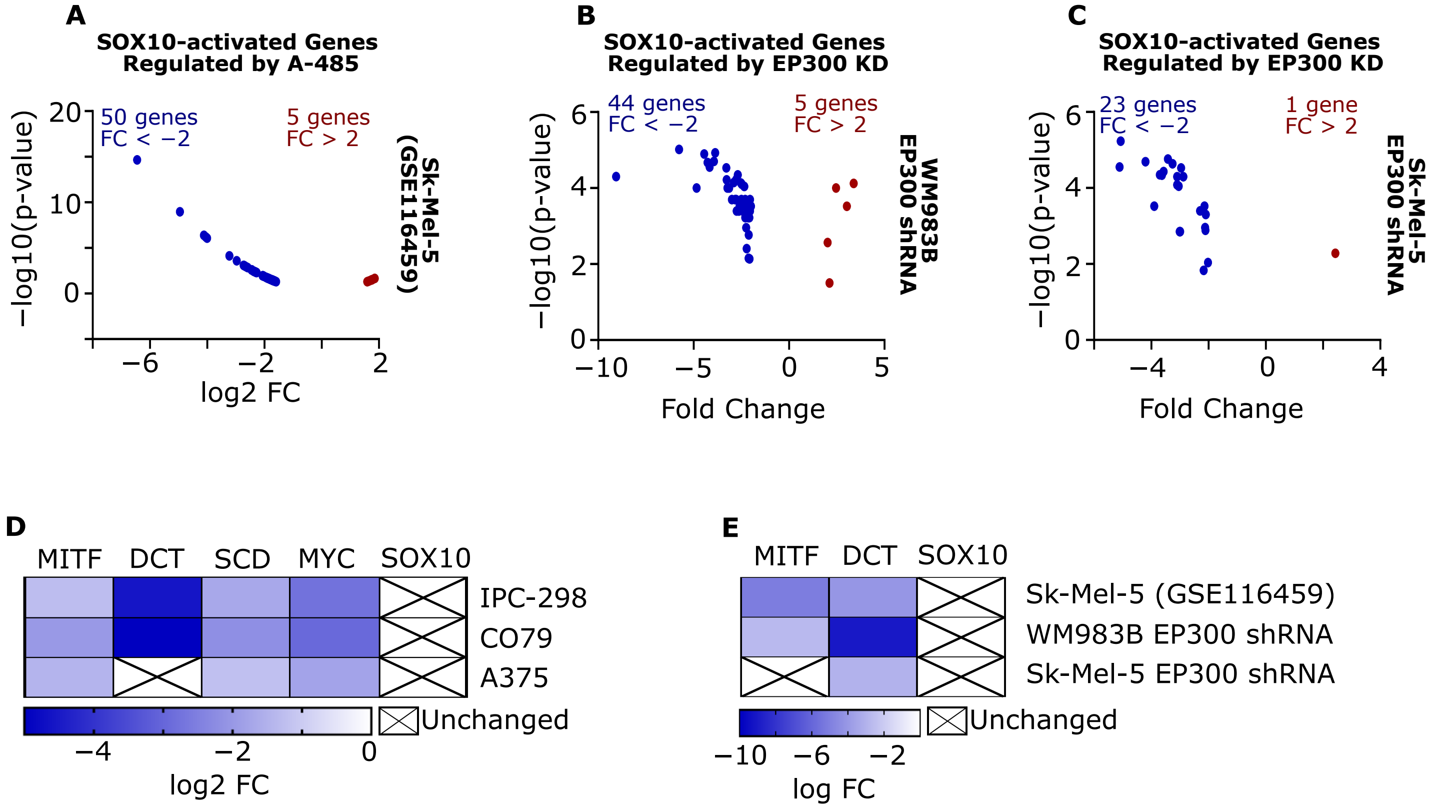

A
SOX10-activated Genes
Regulated by A-485
50 genes
FC < −2
5 genes
FC > 2
Sk-Mel-5
(GSE116459)
−log10(p-value)
20
10
0
−6
−2
2
log2 FC
B
SOX10-activated Genes
Regulated by EP300 KD
44 genes
FC < −2
5 genes
FC > 2
WM983B
EP300 shRNA
−log10(p-value)
6
4
2
0
−10
−5
0
5
Fold Change
C
SOX10-activated Genes
Regulated by EP300 KD
23 genes
FC < −2
1 gene
FC > 2
Sk-Mel-5
EP300 shRNA
−log10(p-value)
6
4
2
0
−4
0
4
Fold Change
D
MITF
DCT
SCD
MYC
SOX10
IPC-298
CO79
A375
Unchanged
−4
−2
0
log2 FC
E
MITF
DCT
SOX10
Sk-Mel-5 (GSE116459)
WM983B EP300 shRNA
Sk-Mel-5 EP300 shRNA
Unchanged
−10
−6
−2
log FC

**Supplementary Figure 5: SOX10-activated genes are reproducibly downregulated by p300 inhibition across multiple datasets. (A)** A volcano plot is shown for genes differentially expressed due to A-485 treatment in Sk-Mel-5 cells from GSE116459. Data analyzed by publicly available GEO2R. **(B)** Volcano plots are shown for genes differentially expressed due to EP300 knockdown in WM983B and Sk-Mel-5 cells (GSE128737). **(C)** SOX10-activated genes downregulated by A-485 in qPCR are reproducibly downregulated in our RNA-seq dataset (see Figure 2). Likewise, SOX10 is not downregulated by qPCR and is also unchanged in our RNA-seq dataset (see Figure 3). **(D)** SOX10 expression is unaltered by A-485 in GSE116459 or EP300 KD in GSE1287373, but SOX10-activated genes are still reproducibly downregulated.

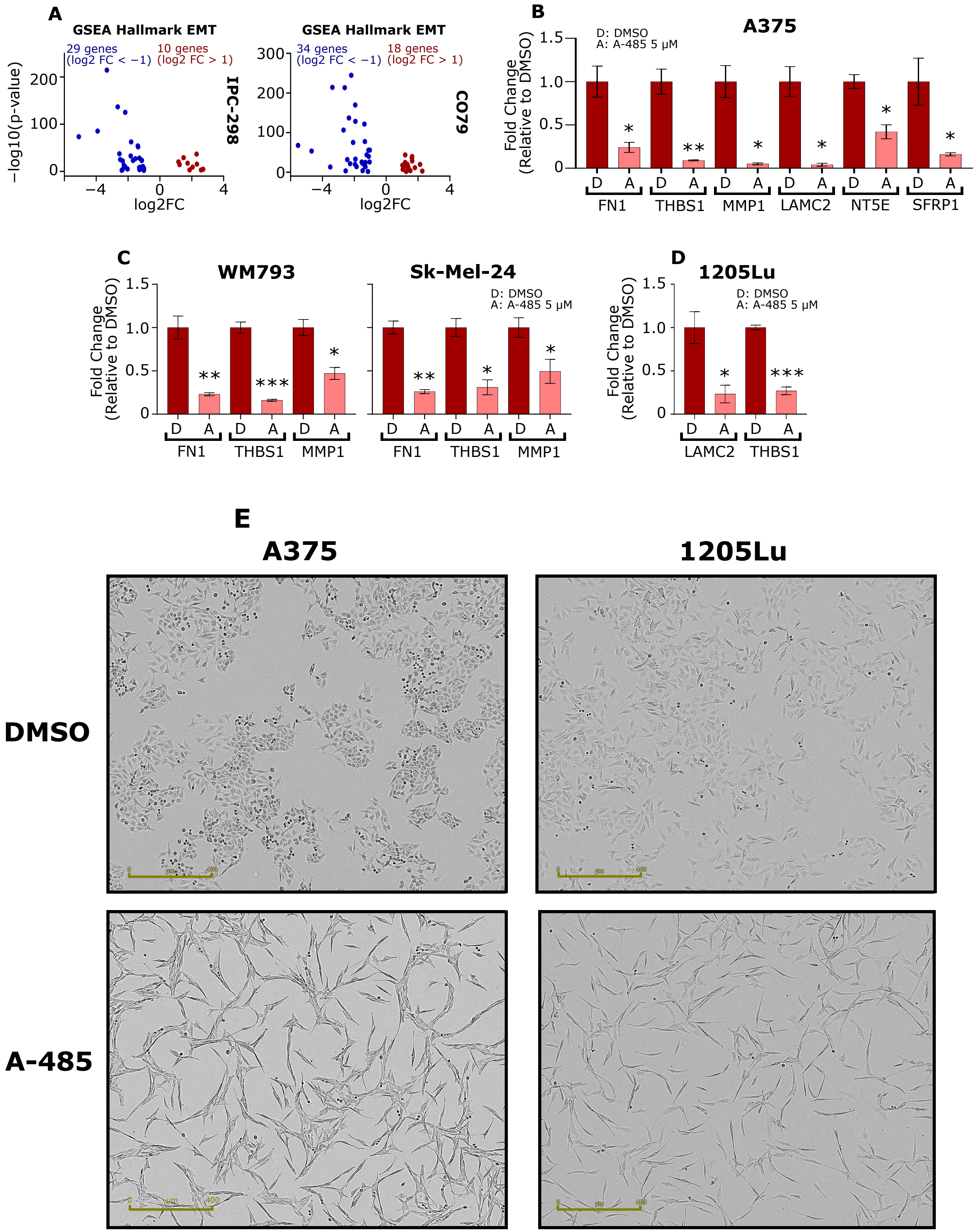
A
GSEA Hallmark EMT
29 genes (log2 FC < −1)
10 genes (log2 FC > 1)
IPC-298
34 genes (log2 FC < −1)
18 genes (log2 FC > 1)
CO79
−log10(p-value)
log2FC
B
A375
D: DMSO
A: A-485 5 μM
Fold Change (Relative to DMSO)
FN1
THBS1
MMP1
LAMC2
NT5E
SFRP1
C
WM793
Sk-Mel-24
D
1205Lu
E
A375
1205Lu
DMSO
A-485

**Supplementary Figure 6: A-485 decreases expression of genes involved in invasion. (A)** Volcano plots are shown for genes in the GSEA EMT Hallmark gene set that are differentially expressed due to A-485 treatment in MITF-high IPC-298 and CO79 cells. **(B)** RT-qPCR validation of SOX10-regulated EMT genes downregulated by A-485 in the RNA-seq in A375 cells. **(C)** RT-qPCR validation of SOX10-regulated EMT genes downregulated by A-485 in WM793 and Sk-Mel-24 cells. **(D)** RT-qPCR validation of SOX10-regulated EMT genes downregulated by A-485 in 1205Lu cells. **(E)** 5 µM A-485 alters the cell morphology of A375 and 1205Lu cells after long-term (12-day) treatment (larger images for comparison to Figure 6J). Data are represented as mean ± SEM. $*p < 0.05$; $**p < 0.005$; $***p < 0.0005$.
